# Linoleic Acid-Lyso PG Axis promoting lipid droplet–mitochondria tethering by stabilizing Noncanonically Mitochondrial PPAR β/δ to Ameliorate Microglial Dysfunction in subarachnoid hemorrhage

**DOI:** 10.64898/2026.08.26.747434

**Authors:** Le-xuan Zou, He Hu, Xunzhi Liu, Junda Shen, Sen Gao, Pengfei Ding, Kexin Tian, Xiaoxuan Dong, Xiaoyan Lu, Ruoyu Liu, Xiaojian Li, Zeng Peng, Xiangxin Chen, Chunhua Hang, Wei Li

**Affiliations:** Department of Neurosurgery, Nanjing Drum Tower Hospital, Affiliated Hospital of Medical School, Nanjing University, Nanjing, Jiangsu 210008, China; Department of Neurosurgery, Nanjing Drum Tower Hospital, Clinical College of Nanjing University of Chinese Medicine, Nanjing, Jiangsu 210008, China; Department of Neurosurgery, Nanjing Drum Tower Hospital, Clinical College of Nanjing Medical University, Nanjing, Jiangsu 210008, China

**Keywords:** subarachnoid hemorrhage, microglia, linoleic acid, lipid droplets, mitochondria, PPARδ

## Abstract

**Background:** Microglial lipid handling and mitochondrial failure contribute to brain injury after subarachnoid hemorrhage (SAH), but the lipid signals coupling these processes remain unclear. We investigated whether linoleic acid (LA) restores microglial homeostasis through lysophosphatidylglycerol 16:0 (LPG[16:0]) and peroxisome proliferator-activated receptor-δ (PPARδ).

**Methods:** Case-control CSF metabolomics included 30 patients with aneurysmal SAH and 10 controls. Mechanisms were examined in a blood-injection mouse model and hemoglobin-exposed primary mouse microglia using targeted lipidomics, RNA sequencing, mitochondrial and phagocytosis assays, pharmacological perturbation, fractionation, coimmunoprecipitation, thermal shift analysis, and structural modeling. Behavioral outcomes were evaluated by open- field, Y-maze, and Morris water-maze testing.

**Results:** CSF LA concentrations were higher in patients with SAH than in controls and discriminated the groups within this cohort (area under the curve, 0.9967 [95% CI, 0.9859–1.000]; P<0.001). LA attenuated inflammatory activation and restored phagocytosis, mitochondrial membrane potential, respiration, and ATP production in hemoglobin-exposed microglia. LA restored PLA2G15-associated LPG(16:0) levels, which phenocopied these effects.

Transcriptomic and inhibitor analyses identified PPARδ as a downstream effector. LPG(16:0) increased PPARδ stability, and fractionation and protease protection identified a PPARδ pool on the cytosolic face of the outer mitochondrial membrane. PPARδ associated with PLIN2 and CPT1A, promoted lipid droplet–mitochondria apposition, and supported fatty acid oxidation. In mice, LA reduced neuroinflammatory injury and partially improved anxiety-related behavior and spatial memory.

**Conclusions:** The LA–LPG(16:0)–PPARδ axis links glycerophospholipid remodeling to organelle coupling and mitochondrial recovery in microglia after SAH. The preventive dosing paradigm requires validation in clinically relevant treatment settings.

## Introduction

Subarachnoid hemorrhage (SAH) is a form of stroke associated with high mortality and disability, accounting for approximately 5%–10% of all strokes. Approximately 85% of nontraumatic SAH cases are caused by rupture of an intracranial aneurysm.^1^Following aneurysm rupture, blood rapidly enters the subarachnoid space, causing an abrupt increase in intracranial pressure and disruption of cerebrospinal fluid circulation, which subsequently lead to marked reductions in cerebral perfusion pressure and cerebral blood flow.^2–4^ Concurrently, erythrocytes and their degradation products, particularly hemoglobin, heme, and iron, exert sustained toxic effects on brain tissue and trigger interconnected pathological processes, including cerebral microcirculatory dysfunction, cerebral edema, oxidative stress, neuroinflammation, and neuronal death.^5–8^ Despite advances in aneurysm treatment and neurocritical care, secondary brain injury after SAH remains a major determinant of patient outcomes, and the functional state of microglia is considered an important factor governing injury progression and tissue repair.^9–12^The American Heart Association/American Stroke Association (AHA/ASA) also characterizes aneurysmal SAH (aSAH) as a highly morbid and often fatal condition.^13^

Microglia are the resident immune cells of the central nervous system and belong to the mononuclear phagocyte system. They maintain brain homeostasis through immune surveillance, inflammatory regulation, and the clearance of damaged tissue and cellular debris.^14^After SAH, hemorrhage-related stimuli can induce maladaptive microglial activation, accompanied by loss of mitochondrial membrane potential, accumulation of reactive oxygen species, and impaired energy production.^15–18^Their phagocytic and damage-clearing capacities are also markedly compromised.^15,19,20^Growing evidence suggests that these phenotypic changes are not driven solely by inflammatory signaling but are closely associated with the reprogramming of microglial metabolism.^17,21,22^In particular, blood cell membranes and damaged tissues impose a substantial lipid burden on microglia. An imbalance among lipid storage, mobilization, and mitochondrial oxidation can lead to abnormal lipid-droplet accumulation, oxidative damage, and impaired phagocytic function.^21,23,24^However, the key signals that drive microglial lipid metabolic dysfunction after SAH and their relationship with mitochondrial impairment remain incompletely understood.

Linoleic acid (LA) is an essential n-6 polyunsaturated fatty acid that is incorporated into membrane phospholipids and serves as a precursor for diverse bioactive oxylipins, thereby linking membrane remodeling to inflammatory signaling and cellular metabolism.^25^ The effects of LA in the injured central nervous system are context dependent. In cerebral ischemia, LA activates microglial peroxisomes and promotes reactive oxygen species scavenging and β-oxidation-dependent anti-inflammatory metabolic reprogramming.^26^ In demyelinating injury, activation of the LA metabolic pathway and supplementation with conjugated linoleic acid (CLA) promote the restoration of microglial lipid homeostasis and the clearance of damaged tissue through peroxisome proliferator-activated receptor-γ (PPARγ).^27^ Conversely, excessive LA can undergo 5-lipoxygenase-dependent peroxidation and generate reactive aldehydes, indicating that its effects depend on dose, subcellular distribution, and metabolic fate.^28^ Recent clinical lipidomic studies have identified LA- associated disturbances in the cerebrospinal fluid fatty acid profile of patients with SAH, as well as transient elevations in plasma LA and several oxylipins.^21,29^ However, whether these changes reflect pathogenic accumulation, compensatory release, or impaired intracellular transport and utilization remains unclear. In particular, How LA remodels glycerophospholipid signaling to coordinate microglial lipid handling with mitochondrial energy production remains to be elucidated.

Accordingly, this study combined cerebrospinal fluid metabolomics from patients with SAH with a mouse model of SAH and hemoglobin-stimulated primary microglia to systematically investigate how LA regulates microglial lipid metabolism and mitochondrial function. Our findings support a mechanistic model in which LA restores PLA2G15-associated LPG(16:0) production, stabilizes mitochondria-associated peroxisome proliferator-activated receptor-δ (PPARδ), and promotes PLIN2–CPT1A-associated lipid droplet–mitochondria coupling and fatty acid β-oxidation, thereby improving microglial mitochondrial function, suppressing inflammatory activation, and restoring phagocytic capacity. This study identifies an LA– LPG(16:0)–PPARδ signaling axis that links fatty acid metabolic dysregulation after SAH to organelle-level metabolic recovery.

## Materials and Methods

### Data availability

The data that support the findings of this study are available from the corresponding authors on reasonable request. All detailed methods are provided in the Supplemental Methods.

### Human Study Design and CSF Metabolomic Analysis

This single-center, hospital-based, unmatched case-control study included 30 patients with aneurysmal subarachnoid hemorrhage and 10 orthopedic control participants recruited at Nanjing Drum Tower Hospital. The principal inclusion criteria and the exclusion criteria were provided in the Supplemental Methods. Demographic and clinical variables, including age, sex, disease severity, sampling time, and relevant comorbidities, were recorded prospectively. Functional outcome at hospital discharge was assessed using the modified Rankin Scale (mRS). For the exploratory outcome analysis, patients with SAH were categorized as having a favorable outcome (mRS 1–2) or an unfavorable outcome (mRS 3–6).

Cerebrospinal fluid (CSF) samples were collected on postoperative day 3 under standardized conditions. Samples were centrifuged at 3000 rpm for 10 minutes at 4°C, and the supernatants were aliquoted and stored at −80°C until analysis. CSF metabolomic profiling was performed using an LC–MS-based platform. Sample preparation, data acquisition, quality-control procedures, metabolite annotation, data preprocessing, and statistical analysis are described in the Supplemental Methods. The corresponding raw and processed data and study metadata were deposited in MetaboLights under accession MTBLS15036.

The study protocol was approved by the Ethics Committee of Nanjing Drum Tower Hospital (approval No. 2022–294–01). Written informed consent was obtained from all participants or their legally authorized representatives. The study was conducted in accordance with the Declaration of Helsinki and is reported following the STROBE recommendations.

### Animal Experiments and SAH Model

Adult male C57BL/6J mice (8–12 weeks old; 20–26 g) were randomly assigned to experimental groups, and investigators responsible for outcome assessment and data analysis were blinded to group allocation. Under isoflurane anesthesia, SAH was induced by injecting 75 μL of freshly collected, nonheparinized arterial blood from a syngeneic donor mouse into the prechiasmatic cistern over 30 seconds. Sham-operated mice underwent identical procedures without blood injection. LA was administered ad libitum as a 1% vol/vol suspension containing 0.3% wt/vol xanthan gum for 14 consecutive days before SAH induction; control mice received xanthan-gum vehicle alone. Detailed housing conditions, surgical procedures, sample-size rationale, exclusion criteria, mortality, and randomization and blinding procedures are provided in the Supplemental Methods. All procedures were approved by the Institutional Animal Care and Use Committee of Nanjing Drum Tower Hospital (No. 2026AE010573) and are reported following the ARRIVE 2.0 guidelines.

### Primary Cell Cultures and Treatments

Primary microglia and astrocytes were obtained from mixed glial cultures prepared from the cerebral cortices of postnatal day 1 C57BL/6J mice. After 14 days in culture, microglia were collected by orbital shaking, and the remaining adherent cells were used as primary astrocytes. Following a 2-day recovery period, cells were exposed to hemoglobin (25 μmol/L) for 24 hours to model SAH-associated injury in vitro. LA, LPG(16:0), and the indicated pharmacological inhibitors were administered as summarized in the figure legends. Detailed isolation, culture, and treatment procedures are provided in the Supplemental Methods. Primary cortical neurons were separately prepared from postnatal day 1 C57BL/6J mice and used for experiments after 10 days in vitro. Detailed culture conditions are provided in the Supplemental Methods.

### Behavioral Assessments

Behavioral testing was performed by investigators blinded to group allocation. Locomotor activity, anxiety-related behavior, and spatial working memory were evaluated on day 14 after SAH using the open-field and Y-maze tests. Spatial learning and reference memory were further evaluated using the Morris water maze beginning on day 16 after SAH. Animal movement was recorded and analyzed using Any-maze. Detailed testing procedures are provided in the Supplemental Methods.

### Molecular and Cellular Analyses

Gene and protein expression were assessed using reverse-transcription quantitative polymerase chain reaction and immunoblotting, respectively. Protein localization and colocalization were evaluated by immunofluorescence microscopy. Cellular reactive oxygen species, lipid peroxidation, and mitochondrial membrane potential were quantified by flow cytometry, whereas mitochondrial respiration and ATP production were assessed using Seahorse analysis and biochemical assays. Coimmunoprecipitation and subcellular fractionation were used to examine protein interactions and localization. Detailed protocols, antibodies, primers, reagents, imaging parameters, and analysis procedures are provided in the Supplemental Methods.

### Transcriptomic and Lipidomic Analyses

RNA sequencing and LC–MS-based lipidomic profiling were performed in primary microglia from the indicated experimental groups. Differential gene-expression analysis was conducted after predefined quality control and normalization, with multiple comparisons controlled using the Benjamini–Hochberg false-discovery-rate procedure, whereas lipid-abundance differences were evaluated using two-sided Student t tests based on nominal P values and were considered exploratory. Functional enrichment analyses were performed using Gene Ontology and pathway databases. Complete sample-processing, sequencing, mass-spectrometry, data-preprocessing, annotation, normalization, differential-analysis, and enrichment workflows are provided in the Supplemental Methods.

### Computational Structural Analyses

Structural models of mouse PPARδ, PLIN2, and CPT1A were obtained from AlphaFold. LPG(16:0) docking to the PPARδ ligand-binding domain was performed using AutoDock Vina, and PPARδ–PLIN2–CPT1A protein-complex docking was performed using HADDOCK3. The selected PPARδ–LPG(16:0) complex was evaluated using a 100-ns molecular-dynamics trajectory in GROMACS, followed by structural-stability, interaction, and MM/GBSA binding-energy analyses. Complete model-preparation, docking, simulation, and analysis parameters are provided in the Supplemental Methods.

### Statistical Analysis

Statistical analyses were performed using GraphPad Prism version 10.6. Data are presented as mean±SD unless otherwise stated. The individual participant was the independent unit for the clinical analyses, and the individual mouse was the independent unit for the in vivo experiments. For cellular experiments, independently prepared cultures or biological samples were treated as independent units. Technical replicates were averaged within each independent experiment before inferential analysis, unless explicitly stated otherwise.

Comparisons between 2 independent groups were performed using a 2-sided unpaired Student t test or an exact 2-sided Mann–Whitney U test, as specified in the figure legends. Comparisons among more than 2 groups were performed using 1-way ANOVA followed by Tukey multiple-comparisons tests. Morris water-maze acquisition data were analyzed using 2-way repeated-measures ANOVA, with treatment group and training day as factors and with evaluation of the group-by-day interaction, followed by Tukey multiple-comparisons tests.

The area under the receiver operating characteristic curve was tested against the null value of 0.5 and is reported with its 95% CI.

RNA-sequencing differential-expression analysis was performed using DESeq2, with multiple comparisons controlled using the Benjamini–Hochberg false-discovery-rate procedure. Genes with an adjusted P value <0.05 and an absolute log2 fold change >1 were considered differentially expressed. Individual lipid abundances were compared using 2-sided Student t tests based on nominal P values. Because these lipidomic comparisons were not selected using multiplicity-adjusted P values, they were considered exploratory; Benjamini– Hochberg-adjusted P values were additionally reported. All tests were 2-sided, and P<0.05 was considered statistically significant.

## Result

### Cerebrospinal fluid metabolomics in patients identifies linoleic acid as a candidate metabolite associated with SAH

To characterize the metabolic alterations associated with SAH in patients, we performed LC– MS-based metabolomic profiling of cerebrospinal fluid (CSF) samples obtained from 30 patients with SAH and 10 control individuals (sFig. 1a). Pathway-level analyses converged on fatty acid metabolism, highlighting the biosynthesis of unsaturated fatty acids, fatty acid biosynthesis and linoleic acid metabolism; Notably, linoleic acid metabolism represented the most prominent pathway in the enrichment tree map (Fig. 1a,b). Consistent with these pathway-level alterations, CSF linoleic acid concentrations were markedly elevated in patients with SAH compared with controls (P < 0.0001; Fig. 1c), although substantial interindividual heterogeneity was observed within the SAH group. Receiver operating characteristic analysis further showed that CSF linoleic acid effectively distinguished patients with SAH from controls within this cohort (AUC = [0.9967], 95% CI = [ 0.9859-1.000]; Fig. 1d). Within the SAH cohort, CSF LA concentrations were numerically higher in patients with favorable discharge outcomes (mRS 1–2) than in those with unfavorable outcomes (mRS 3– 6), although the difference did not reach statistical significance (P=0.0555; sFig. 1b).

**Figure 1.**
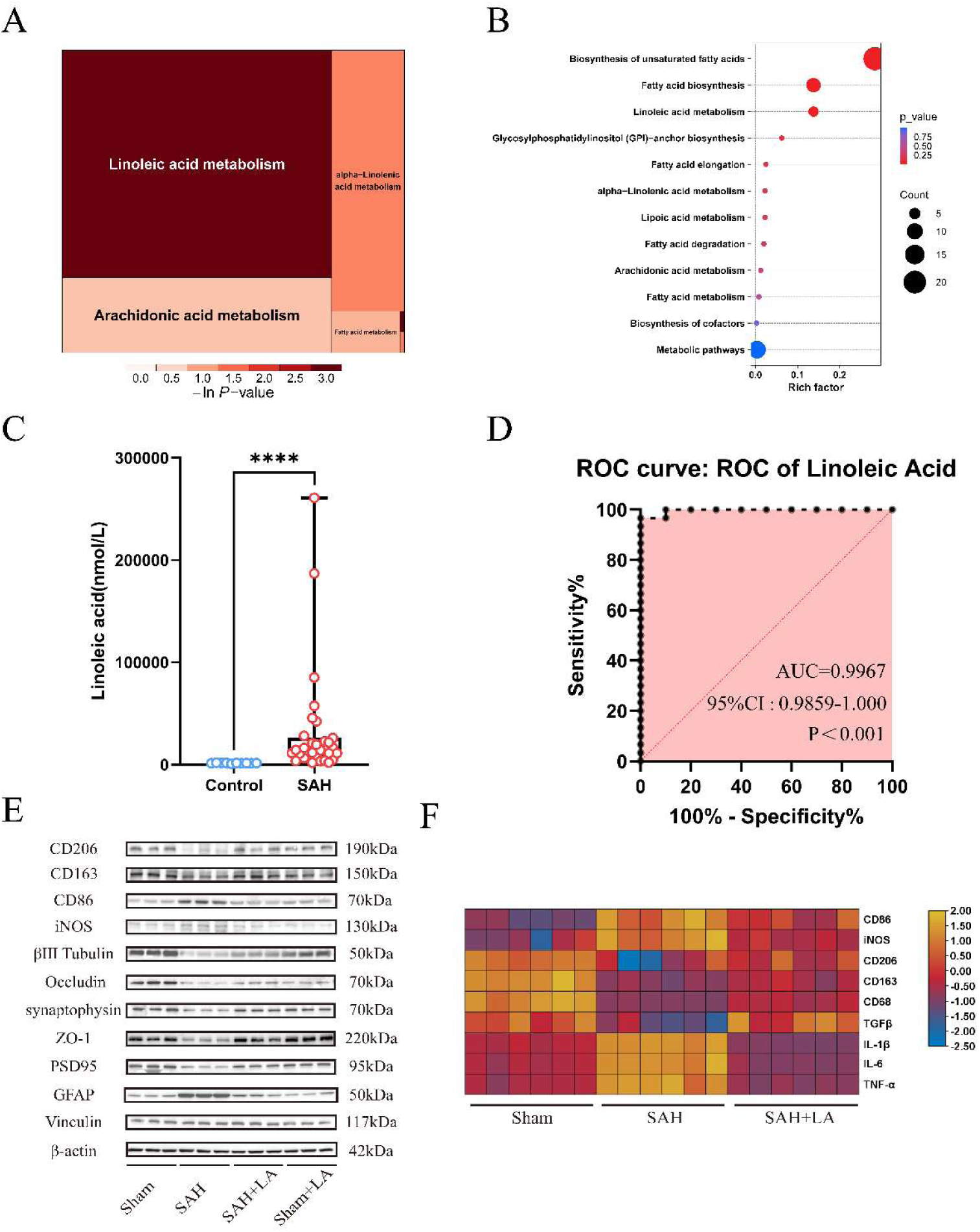
Cerebrospinal fluid metabolomics identifies linoleic acid dysregulation in subarachnoid hemorrhage, and linoleic acid attenuates neuroinflammatory injury in mice. **A**, Treemap of metabolic pathways altered in cerebrospinal fluid (CSF) from patients with subarachnoid hemorrhage (SAH) relative to control participants; tile size and color represent pathway enrichment and -ln(P value), respectively. **B**, Kyoto Encyclopedia of Genes and Genomes pathway-enrichment bubble plot showing the rich factor, number of annotated metabolites, and P value for the indicated pathways. **C**, CSF linoleic acid concentrations in control participants (n=10) and patients with SAH (n=30). **D**, Receiver operating characteristic curve for CSF linoleic acid discrimination of SAH in this cohort (area under the curve, 0.9967; 95% CI, 0.9859-1.000; P<0.001). **E**, Representative immunoblots of microglial activation markers (CD206, CD163, CD86, and inducible nitric oxide synthase [iNOS]), neuronal/synaptic proteins (βIII-tubulin, synaptophysin, and PSD95), blood-brain barrier proteins (occludin and ZO-1), and the astrocytic marker GFAP in temporal-lobe tissue from sham, SAH, SAH+linoleic acid (LA), and sham+LA mice. Three representative animals per group are shown; the corresponding densitometric analysis is presented in Supplemental Figure 1D, and the complete animal cohort comprised n=6 per group. **F**, Heat map of standardized Il1b, Il6, Tnf, Tgfb, Cd163, Cd206, Nos2, and Cd86 mRNA expression in temporal-lobe tissue from sham, SAH, and SAH+LA mice (n=6 per group). LA was provided as a 1% vol/vol suspension in 0.3% wt/vol xanthan gum for 14 days before SAH induction. Data are mean±SD. Data in C were compared using a 2-sided unpaired Student t test; ****P<0.0001. Multi-group mRNA data in F were analyzed using 1-way ANOVA followed by the Tukey multiple-comparisons test. The ROC P value tests the null hypothesis that the area under the curve equals 0.5. Immunoblot band intensities were quantified using Fiji/ImageJ (version 1.54p). AUC indicates area under the curve; CI, confidence interval; CSF, cerebrospinal fluid; GFAP, glial fibrillary acidic protein; LA, linoleic acid; ROC, receiver operating characteristic; and SAH, subarachnoid hemorrhage.

Collectively, these clinical findings identify elevated CSF linoleic acid and perturbation of linoleic acid-associated metabolism as prominent metabolic features of SAH, providing the rationale for subsequent mechanistic investigation.

### LA treatment attenuates neurological injury and neuroinflammation after subarachnoid hemorrhage

To characterize the therapeutic potential of Linoleic Acid (LA) after subarachnoid hemorrhage (SAH), we examined the expression of inflammatory cytokine genes post SAH in vivo, as neuroinflammation is a central pathological process in early brain injury after SAH using real-time quantitative reverse transcription PCR (RT-PCR). The result shows that LA treatment significantly suppressed SAH-induced upregulation of IL-1β, IL-6, and TNF-α at transcriptional level (Fig. 1f). Consistent with these anti-inflammatory effects, Glial cells as the central regulators of the pathological response to SAH stimulation, LA treatment attenuated astrocyte and microglial activation after SAH (Fig. 1e; sFig. 1d). These effects were accompanied by restored expression of synaptic proteins (βIII-tubulin and synaptophysin) and tight junction proteins (occludin and ZO-1) (Fig. 1e; sFig. 1d), showing the protective effect of LA treatment in neuronal network after SAH. Taken together, LA treatment exhibited broadly involved therapeutic effect in Neuron, Astrocytes and Microglia within the pathological environment after SAH.

### LA selectively restores mitochondrial function in microglia after subarachnoid hemorrhage

After observing that LA reduced neuroinflammation and tissue injury in vivo, we next sought to identify the cell type directly responsive to LA. Referring to the pleiotropic effect of LA in SAH, Primary microglia, Primary astrocytes, and Primary neurons were cultured separately and exposed to a SAH-mimicking insult, followed by LA treatment. We assessed microglial activation by measuring CD86, iNOS, CD206 and CD163, astrocytic activation by GFAP, and neuronal injury by synaptic proteins, including βIII-tubulin, synaptophysin and PSD95 (Fig. 2a; sFig. 2a). Notably, LA markedly attenuated activation-associated changes in microglia but did not directly reduce astrocytic activation or neuronal injury in monoculture (Fig. 2a; sFig. 2a). These findings identified microglia as the principal LA-responsive cell population under these in vitro conditions.

**Figure 2.**
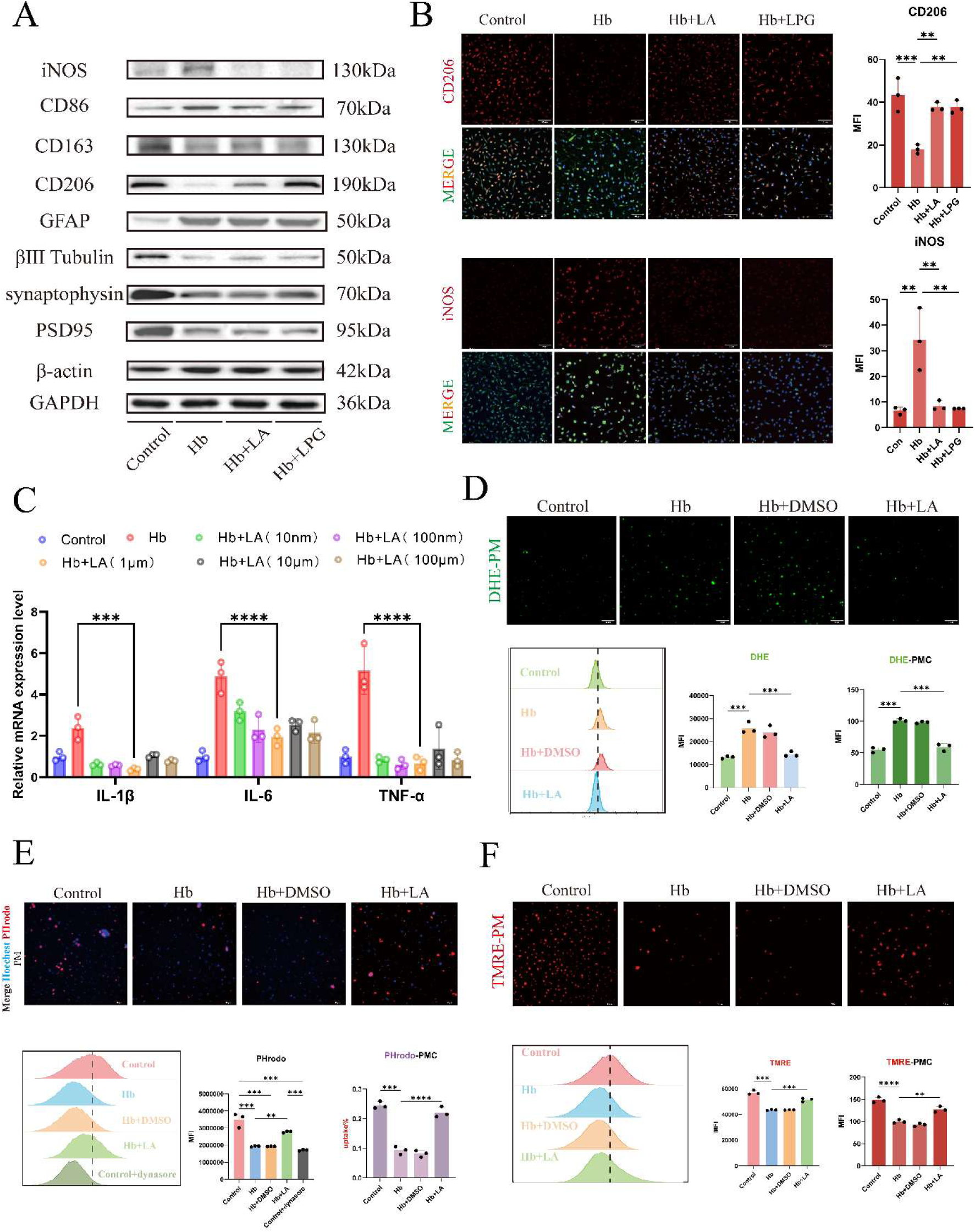
Linoleic acid and lysophosphatidylglycerol 16:0 suppress hemoglobin-induced microglial activation and restore microglial function. Primary mouse microglia were exposed to hemoglobin (Hb; 25 μmol/L) for 24 hours and treated with linoleic acid (LA; 1 μmol/L) or lysophosphatidylglycerol 16:0 (LPG[16:0]; 50 nmol/L), unless otherwise indicated. **A**, Representative immunoblots of microglial activation markers (iNOS, CD86, CD163, and CD206), the astrocytic marker GFAP, and neuronal/synaptic proteins (βIII-tubulin, synaptophysin, and PSD95) in primary microglia, astrocytes, and cortical neurons under the indicated conditions; the corresponding densitometric analysis is presented in Supplemental Figure 2A. **B**, Representative immunofluorescence images and mean fluorescence intensity (MFI) quantification of CD206 and iNOS in primary microglia. **C**, Relative Il1b, Il6, and Tnf mRNA expression after treatment with increasing concentrations of LA (10 nmol/L–100 μmol/L). **D**, Representative dihydroethidium (DHE) fluorescence images, flow-cytometry distributions, and MFI quantification of oxidant-associated fluorescence in Hb-exposed primary microglia treated with dimethyl sulfoxide vehicle or LA. **E**, Representative pHrodo Red E. coli BioParticle uptake images, flow-cytometry distributions, MFI, and pHrodo-positive fraction. Dynasore (40 μmol/L) served as a phagocytosis inhibitor. **F**, Representative tetramethylrhodamine ethyl ester (TMRE) fluorescence images, flow-cytometry distributions, and MFI quantification of mitochondrial membrane potential. Scale bars are shown in the images. Data are mean±SD from 3 independent cultures. Comparisons were performed using 1-way ANOVA followed by Tukey multiple-comparisons tests; *P<0.05, **P<0.01, ***P<0.001, and ****P<0.0001. Fluorescence images were acquired with the Leica THUNDER imaging system and processed using LAS X version 3.7.22383.2 and Fiji/ImageJ version 1.54p; flow-cytometry data were analyzed using FlowJo version 10.10.0. DHE indicates dihydroethidium; DMSO, dimethyl sulfoxide; GFAP, glial fibrillary acidic protein; Hb, hemoglobin; iNOS, inducible nitric oxide synthase; LA, linoleic acid; LPG(16:0), lysophosphatidylglycerol 16:0; MFI, mean fluorescence intensity; and TMRE, tetramethylrhodamine ethyl ester.

Having established microglia as a major cellular target of LA, we next investigated how LA modulated microglial activation after SAH. Because microglia are resident immune cells of the central nervous system and belong to the mononuclear phagocyte lineage, we examined both inflammatory signaling and phagocytic function. LA treatment significantly reduced Hb-induced transcription of inflammatory cytokines in microglia and IF result showed that LA treatment attenuated activation in microglia (Fig. 2b,c). In parallel, pHrodo-based phagocytosis assays showed that LA restored the impaired phagocytic activity of microglia after SAH, as reflected by increased mean fluorescence intensity (Fig,2e).

We then asked whether these anti-inflammatory and pro-phagocytic effects were associated with changes in oxidative stress and mitochondrial function. Flow cytometry and DHE staining showed that LA significantly reduced intracellular ROS accumulation in microglia after SAH (Fig. 2d). Because excessive ROS generation and impaired phagocytosis are closely linked to mitochondrial dysfunction, we further assessed mitochondrial activity.

TMRE and JC-1 assays showed that LA restored mitochondrial membrane potential (Fig. 2f,3f), while ATP measurements and Seahorse XF Pro analysis demonstrated improved ATP production and oxygen consumption rate (OCR) after LA treatment (Fig. 7a,7b).

Together, these results indicate that LA attenuated inflammatory activation and restored phagocytic function in microglia, at least in part by preserving mitochondrial function after SAH.

### Targeted lipidomic and phenotypic validation identify PLA2G15-associated LPG(16:0) restoration as a key mechanism of LA action

Having shown that LA restored mitochondrial function, improved phagocytic activity, and reduced inflammatory activation in microglia after SAH, we next sought to define the downstream lipid mediators associated with these effects. Given that LA is an essential polyunsaturated fatty acid, we performed targeted lipidomic profiling to quantify LA-induced changes in the lipid pool of microglia after SAH. Principal component analysis showed clear separation among the experimental groups, supporting broad lipidomic remodeling after SAH and LA treatment (sFig. 3a). A total of 376 metabolites were commonly detected across groups, and KEGG pathway analysis indicated prominent enrichment in glycerophospholipid metabolism (Fig. 3a).

**Figure 3.**
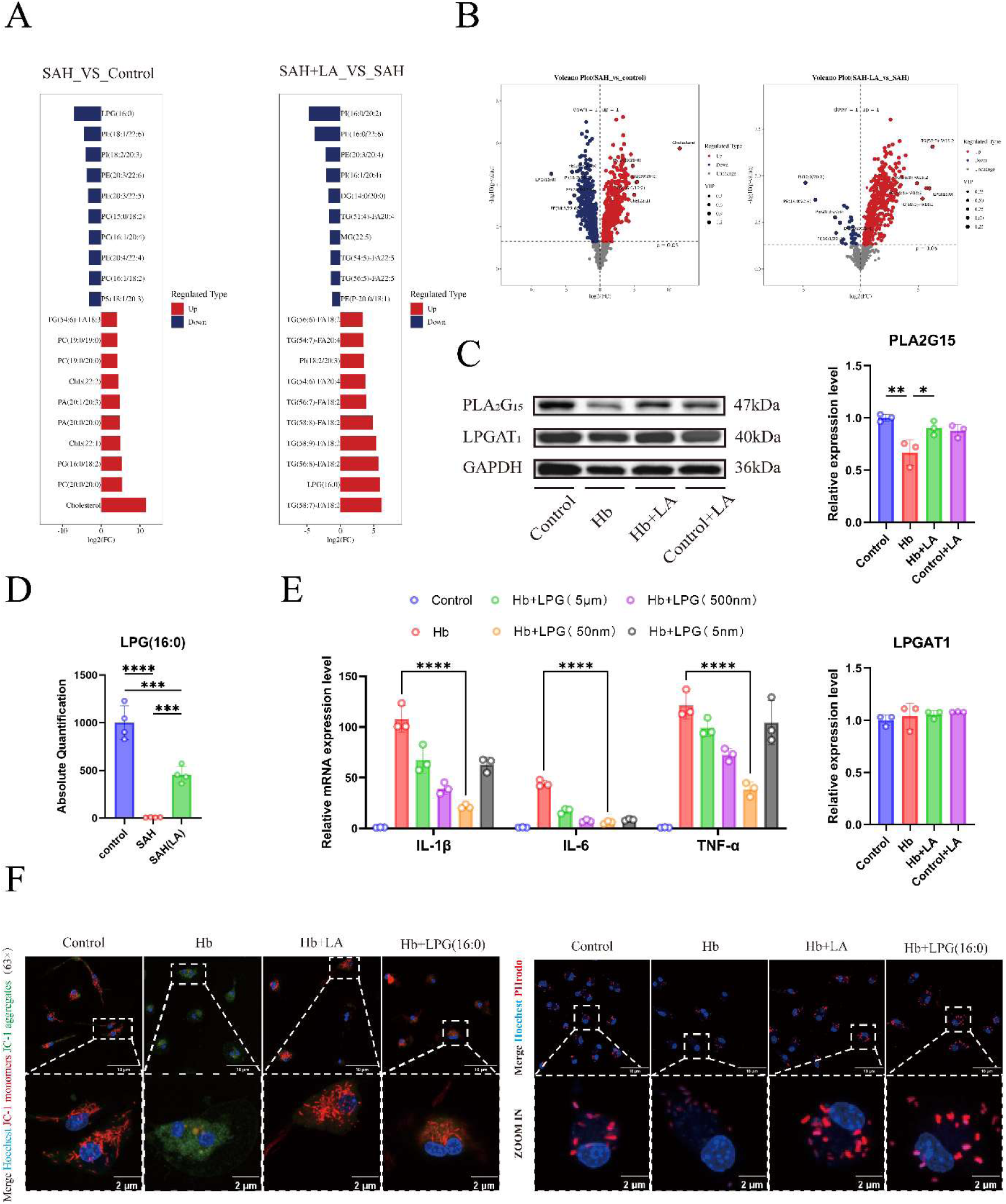
Linoleic acid restores PLA2G15-associated lysophosphatidylglycerol 16:0 abundance in hemoglobin-exposed primary microglia. **A**, Differentially abundant lipid species in SAH versus control and SAH+linoleic acid (LA) versus SAH comparisons, ranked by log2 fold change. **B**, Volcano plots of lipid-abundance changes for the corresponding comparisons; color indicates the direction of change, and point size represents variable importance in projection. Targeted lipidomics included 4 biological samples per group. **C**, Representative immunoblots and densitometric quantification of PLA2G15 and LPGAT1 in primary microglia exposed to hemoglobin (Hb; 25 μmol/L for 24 hours) with or without LA (1 μmol/L). **D**, Absolute LPG(16:0) abundance measured by targeted lipidomics in control, SAH, and SAH+LA groups (n=4 biological samples per group). **E**, Relative Il1b, Il6, and Tnf mRNA expression in Hb-exposed primary microglia treated with LPG(16:0) at 5 nmol/L, 50 nmol/L, 500 nmol/L, or 5 μmol/L. **F**, Representative JC-1 fluorescence images showing mitochondrial membrane potential and representative pHrodo Red E. coli BioParticle uptake images in control and Hb-exposed primary microglia treated with LA or LPG(16:0) (50 nmol/L). Scale bars are shown in the images. Cell- validation data are mean±SD from 3 independent cultures and were analyzed using 1-way ANOVA followed by Tukey multiple-comparisons tests; *P<0.05, **P<0.01, ***P<0.001, and ****P<0.0001. Individual lipid abundances were screened using 2-sided Student t tests with nominal P<0.05; these analyses were exploratory, with fold changes, variable- importance-in-projection scores, and Benjamini-Hochberg false-discovery-rate-adjusted P values additionally reported. Fluorescence images were acquired with the Leica STELLARIS 5 confocal microscope. Immunoblots and fluorescence images were quantified using Fiji/ImageJ version 1.54p. Hb indicates hemoglobin; JC-1, 5,5′,6,6′-tetrachloro-1,1′,3,3′- tetraethylbenzimidazolylcarbocyanine iodide; LA, linoleic acid; LPG(16:0), lysophosphatidylglycerol 16:0; LPGAT1, lysophosphatidylglycerol acyltransferase 1; PLA2G15, phospholipase A2 group XV; SAH, subarachnoid hemorrhage; and VIP, variable importance in projection.

Among the altered lipid metabolites, 1-palmitoyl-2-hydroxy-*sn*-glycero-3-phospho-(1′-*rac*- glycerol) [LPG(16:0)] showed one of the most pronounced LA-responsive changes. SAH caused an approximately 136-fold depletion of LPG(16:0), whereas LA treatment increased LPG(16:0) by nearly 53-fold relative to the SAH group, partially restoring the nearly exhausted LPG(16:0) pool (Fig. 3d). This selective restoration prompted us to determine whether LPG(16:0) itself could reproduce the protective effects of LA in microglia. The result shows that treatment with LPG(16:0) largely phenocopied the effects of LA after SAH, restoring mitochondrial membrane potential and OCR, reducing excessive ROS accumulation, suppressing inflammatory cytokine production, and improving impaired phagocytic function (Fig. 3f). These findings identify LPG(16:0) as a key downstream lipid mediator of LA action in microglia.

We next investigated how LA restored LPG(16:0) abundance after SAH. PLA2G15 and LPGAT1 were selected as candidate enzymes involved in LPG metabolism, representing putative synthetic and degradative arms of this pathway, respectively. PLA2G15 expression was markedly reduced in microglia after SAH, consistent with the depletion of LPG species observed in the lipidomic analysis. LA treatment reversed this SAH-induced reduction in PLA2G15 expression (Fig. 3c). By contrast, LPGAT1 expression was not significantly altered after SAH or LA treatment (Fig. 3c). Together, these data suggest that LA restores the depleted LPG(16:0) pool primarily by reversing SAH-induced PLA2G15 downregulation, thereby preserving mitochondrial function and microglial homeostasis after SAH.

### Transcriptomics and inhibitor screening identify PPARδ as a key downstream target of the LA–LPG axis

To further define the mechanism by which the LA–LPG axis restored microglial function, we performed transcriptomic profiling of SAH-modeled microglia treated with LA or LPG(16:0).

Principal component analysis showed clear separation among the experimental groups, supporting distinct transcriptional remodeling after LA and LPG(16:0) treatment (sFig. 4b). Venn analysis identified 68 overlapping genes regulated by both LA and LPG(16:0) (sFig. 4c). KEGG pathway enrichment analysis of these shared genes revealed significant enrichment of the PPAR signaling pathway, and concordantly regulated genes in the nine-quadrant analysis further supported activation of this pathway (Fig. 4a).

**Figure 4.**
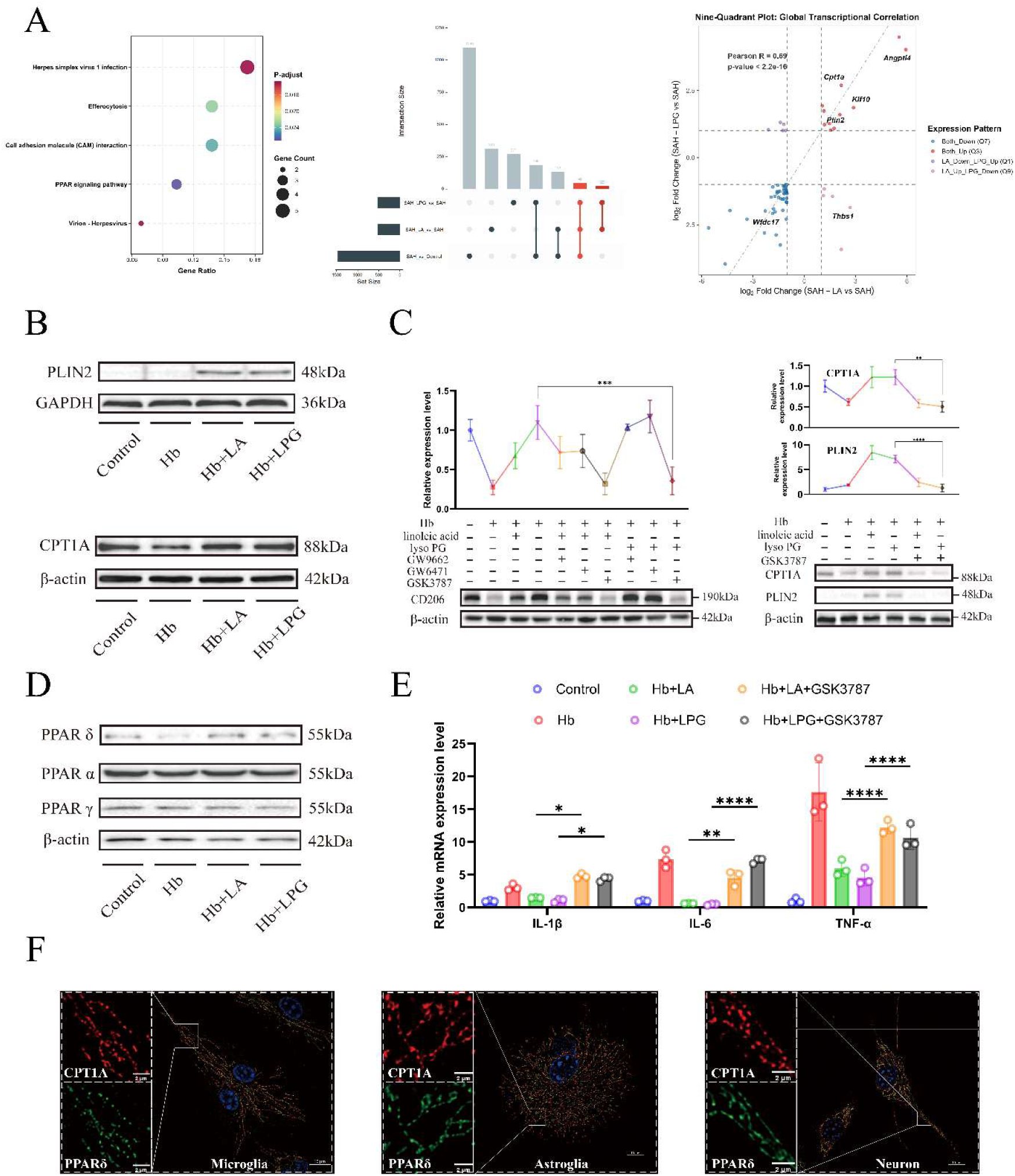
Transcriptomic and pharmacological analyses identify PPARδ as a downstream effector of the linoleic acid–lysophosphatidylglycerol axis. **A**, Kyoto Encyclopedia of Genes and Genomes enrichment analysis of genes jointly regulated by linoleic acid (LA) and lysophosphatidylglycerol 16:0 (LPG[16:0]), UpSet plot of differentially expressed gene intersections, and 9-quadrant plot comparing transcriptional changes induced by LA and LPG(16:0) in hemoglobin (Hb)-exposed primary microglia. RNA sequencing included 4 biological samples per group. Differential expression was defined as a Benjamini-Hochberg-adjusted P<0.05 and an absolute log2 fold change >1. **B**, Representative immunoblots of PLIN2 and CPT1A in control and Hb-exposed primary microglia treated with LA or LPG(16:0). **C**, PPAR isoform inhibitor screening using CD206 expression and validation of the effects of the PPARδ antagonist GSK3787 on CPT1A and PLIN2. GW9662 (10 μmol/L), GW6471 (5 μmol/L), and GSK3787 (10 μmol/L) were used to inhibit PPARγ, PPARα, and PPARδ, respectively. **D**, Representative immunoblots of PPARδ, PPARα, and PPARγ. **E**, Relative Il1b, Il6, and Tnf mRNA expression after GSK3787 treatment in Hb- exposed primary microglia treated with LA or LPG(16:0). **F**, Representative super-resolution immunofluorescence images showing PPARδ and CPT1A localization in primary microglia, astrocytes, and cortical neurons. Scale bars are shown in the images. Cell-validation data are mean±SD from 3 independent cultures and were analyzed using 1-way ANOVA followed by Tukey multiple-comparisons tests; *P<0.05, **P<0.01, ***P<0.001, and ****P<0.0001. Immunoblot signals were quantified using Fiji/ImageJ version 1.54p; super-resolution images were acquired using the ZEISS Lattice SIM 5 and processed with ZEN version 3.9. CPT1A indicates carnitine palmitoyltransferase 1A; Hb, hemoglobin; LA, linoleic acid; LPG(16:0), lysophosphatidylglycerol 16:0; PLIN2, perilipin 2; and PPAR, peroxisome proliferator- activated receptor.

Because the Peroxisome Proliferator-Activated Receptors (PPARs) family comprises three major lipid metabolism-associated nuclear receptors, PPARα, PPARβ/δ, and PPARγ, transcriptomic enrichment alone could not determine which receptor mediated the LA–LPG response. We therefore performed a functional inhibitor screen using CD206 as a phenotypic marker of microglial reparative activation. The PPARδ-specific inhibitor GSK3787 abolished the LA- and LPG(16:0)-induced restoration of CD206 expression, whereas inhibition of PPARα or PPARγ did not significantly affect this response (Fig. 4c). In parallel, GSK3787 reversed the anti-inflammatory effect of the LA–LPG axis, increasing inflammatory cytokine expression in SAH-modeled microglia (Fig. 4e). Moreover, canonical downstream proteins of PPARδ that were restored by LA or LPG(16:0) were reduced to levels comparable to the SAH group after GSK3787 treatment (Fig. 4c). Besides, Among the three PPAR isoforms examined, only PPARδ showed a significant change in protein expression in the SAH model, which was restored by LA-LPG treatment (Fig. 4d).

Together, these transcriptomic and pharmacological validation experiments identify PPARδ as a key downstream effector of the LA–LPG axis in microglia after SAH.

### A mitochondria-associated pool of PPARδ localizes to the cytosolic face of the outer **mitochondrial membrane in microglia**

As PPARδ is classically recognized as a nuclear receptor, we first examined its subcellular distribution in microglia after SAH by immunofluorescence staining. Unexpectedly, super-resolution microscopy revealed that PPARδ did not display a predominantly diffuse nuclear pattern. Instead, PPARδ immunoreactivity appeared as elongated, filament-like structures and showed marked colocalization with CPT1A, an outer mitochondrial membrane protein (Fig. 4f). Consistently, subcellular fractionation followed by western blot analysis showed that PPARδ was enriched mainly in the mitochondrial fraction, rather than being restricted to the nuclear compartment (Fig. 5a). These results indicate that, in addition to its canonical nuclear localization, PPARδ has a non-canonical mitochondria-associated pool in microglia.

**Figure 5.**
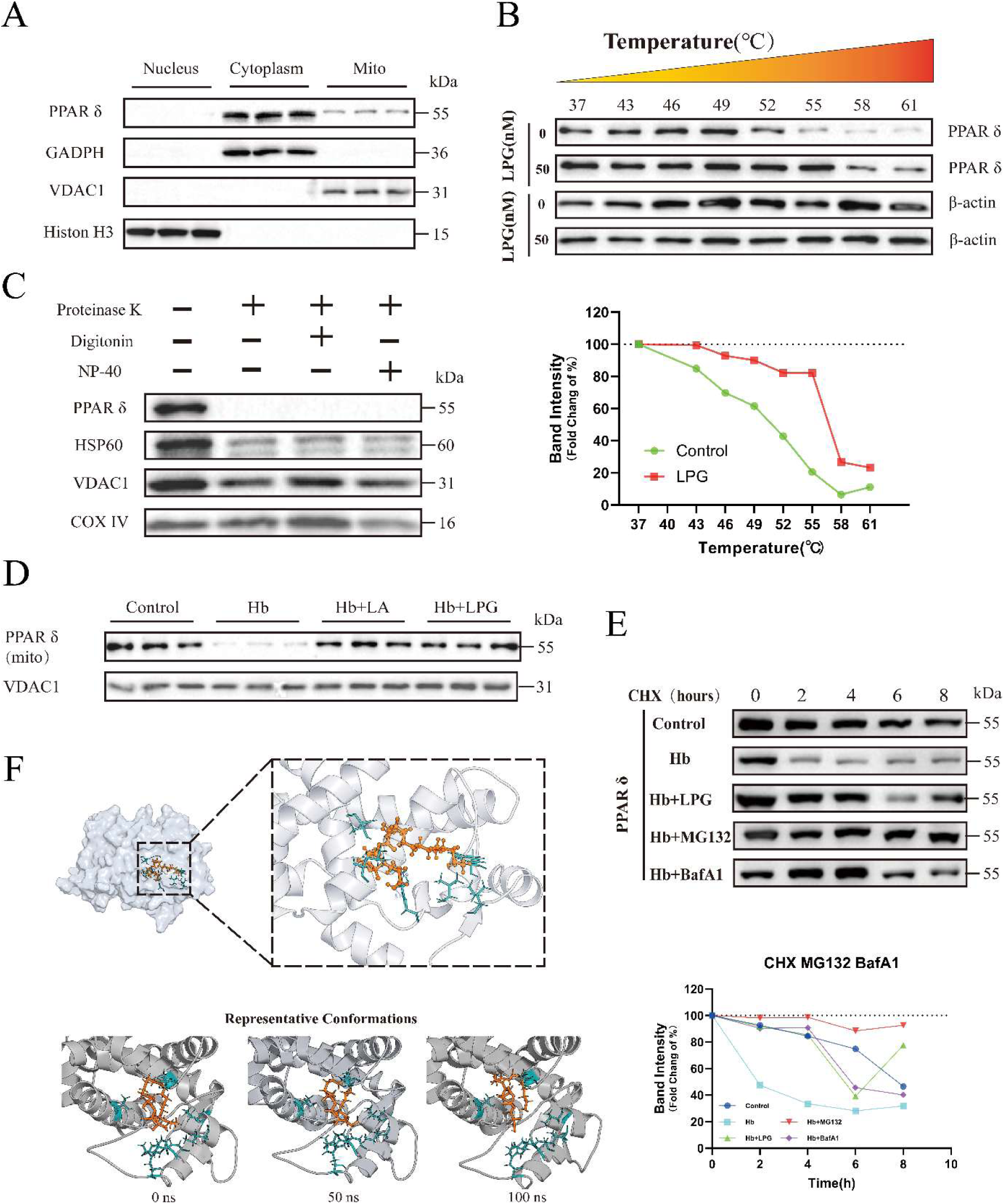
Lysophosphatidylglycerol 16:0 stabilizes a mitochondria-associated pool of PPARδ. **A**, Subcellular fractionation and immunoblot analysis of PPARδ in nuclear, cytoplasmic, and mitochondrial fractions of primary microglia. Histone H3, GAPDH, and VDAC1 served as nuclear, cytoplasmic, and mitochondrial markers, respectively. **B**, Cellular thermal shift assay of PPARδ in the absence or presence of lysophosphatidylglycerol 16:0 (LPG[16:0]; 50 nmol/L) across the indicated temperatures, with normalized band-intensity curves. **C**, Protease-protection analysis of isolated mitochondria treated with proteinase K (20 μg/mL), digitonin (0.1 mg/mL), or NP-40 (1% vol/vol). HSP60, VDAC1, and COX IV served as mitochondrial matrix, outer-membrane, and inner-membrane markers, respectively. **D**, Representative immunoblots of mitochondria-associated PPARδ in control and hemoglobin (Hb)-exposed primary microglia treated with linoleic acid (LA; 1 μmol/L) or LPG(16:0). **E**, Cycloheximide chase analysis of PPARδ degradation at 0, 2, 4, 6, and 8 hours after inhibition of protein synthesis with cycloheximide (10 μg/mL), with LPG(16:0), MG132 (10 μmol/L), or bafilomycin A1 (50 nmol/L) treatment as indicated; curves show normalized band intensity. **F,** Predicted binding pose of LPG(16:0) within the mouse PPARδ ligand-binding domain and representative conformations at 0, 50, and 100 ns during molecular-dynamics simulation. Immunoblot experiments were independently repeated 3 times; representative results are shown. Band intensities were quantified using Fiji/ImageJ version 1.54p. COX IV indicates cytochrome c oxidase subunit IV; GAPDH, glyceraldehyde-3-phosphate dehydrogenase; Hb, hemoglobin; HSP60, heat shock protein 60; LA, linoleic acid; LPG(16:0), lysophosphatidylglycerol 16:0; PPARδ, peroxisome proliferator-activated receptor δ; and VDAC1, voltage-dependent anion channel 1.

Furthermore, immunofluorescence analyses in neurons and astrocytes supported the mitochondria-associated localization of PPARδ, suggesting that this subcellular localization pattern is conserved across major brain cell types (Fig. 4f).

Because mitochondrial localization can correspond to distinct submitochondrial compartments and functions, we next determined the precise topology of mitochondrial PPARδ. Isolated intact mitochondria were sequentially treated with proteinase K, digitonin and NP-40 to progressively expose mitochondrial membrane compartments. PPARδ was rapidly lost after proteinase K digestion of intact mitochondria, whereas proteins protected within inner mitochondrial compartments were retained under the same condition (Fig. 5c). This protease sensitivity indicates that mitochondria-associated PPARδ is exposed on the cytosolic face of the outer mitochondrial membrane, without a detectable membrane-protected domain. Together, these findings identify a non-canonical outer mitochondrial membrane-associated localization of PPARδ in microglia, suggesting that PPARδ may exert mitochondria-related functions beyond its established role as a nuclear receptor.

### LPG stabilizes mitochondria-associated PPARδ by suppressing ubiquitin and lysosome-dependent degradation

PPARδ is a rapidly turning-over protein, and we found that the mitochondria-associated pool of PPARδ was more prominently reduced than total cellular PPARδ in SAH microglia (Fig. 5d). We therefore asked whether the SAH-related injury environment accelerates PPARδ degradation and whether LPG preserves mitochondrial PPARδ by stabilizing the protein. In cycloheximide chase assays, Hb stimulation markedly accelerated the loss of PPARδ protein, whereas LPG treatment slowed PPARδ degradation (Fig. 5e), indicating that LPG maintains PPARδ expression at least in part by increasing protein stability. Consistently, cellular thermal shift assay analysis showed that LPG restored the thermal stability of PPARδ, supporting a stabilizing effect of LPG on PPARδ (Fig. 5b).

We next examined whether this stabilization could involve a direct LPG–PPARδ interaction. In 100-ns molecular dynamics simulations, the LPG(16:0)–PPARδ complex remained globally stable, as indicated by stable RMSD, Cα RMSF, solvent-accessible surface area and radius of gyration profiles (sFig. 5b,c,d). Although LPG showed expected conformational flexibility as a lipid ligand, it remained within a similar binding region throughout the simulation and did not undergo obvious dissociation at 0, 50 or 100 ns (Fig. 5f). Hydrogen-bond analysis showed intermittent but recurrent polar interactions between LPG and PPARδ, while electrostatic surface analysis suggested that the polar head group of LPG was positioned near a charged surface environment (sFig. 5e,l). Free-energy landscape analysis identified several low-energy basins, indicating that the complex sampled stable or metastable conformational states rather than undergoing random conformational drift (sFig. 5k). MM/GBSA analysis further predicted a favorable binding energy for the LPG– PPARδ complex, with an average ΔG bind of approximately −52.43 kcal/mol from 20 to 100 ns (sFig. 5g). Energy decomposition indicated that van der Waals and nonpolar hydrophobic interactions were the major contributors, with Leu293, Asn306, Cys248, Met191, Ile296, Ile289 and Thr251 contributing prominently to ligand binding (sFig. 5i). Together, these simulations support a dynamic but persistent interaction between LPG(16:0) and PPARδ.

To define the degradation route responsible for PPARδ loss, we inhibited protein synthesis with cycloheximide and then blocked proteasomal or lysosomal degradation using MG132 or BafA1, respectively. Both inhibitors reduced the Hb-induced acceleration of PPARδ degradation (Fig. 5e), suggesting that PPARδ is controlled by both proteasome- and lysosome-related degradation pathways under SAH-like stress. LPG similarly suppressed the Hb-induced increase in PPARδ degradation. Western blot analysis of global ubiquitination, together with immunoprecipitation of PPARδ followed by ubiquitin detection, further showed that Hb increased PPARδ ubiquitination, whereas LPG reduced this effect (Fig. 6b). In parallel, analysis of lysosome-related functional proteins supported the involvement of lysosomal degradation in Hb-induced PPARδ loss (Fig. 6c,d). These data indicate that LPG directly stabilizes PPARδ and protects the mitochondria-associated PPARδ pool by limiting Hb-induced ubiquitin- and lysosome-dependent degradation.

**Figure 6.**
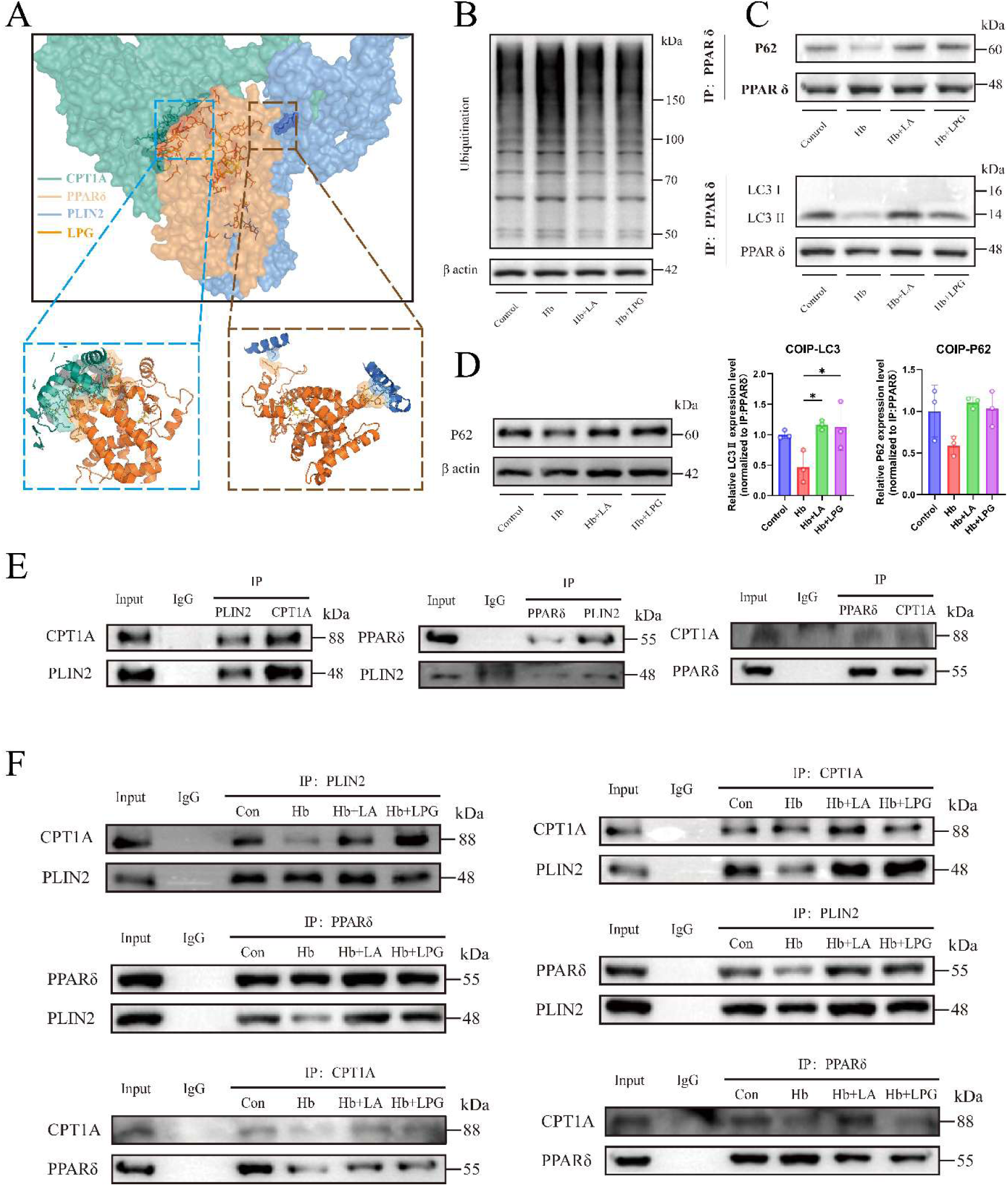
PPARδ forms a complex with PLIN2 and CPT1A and is protected from ubiquitin- and autophagy-associated degradation by the linoleic acid– lysophosphatidylglycerol axis. **A**, HADDOCK3 model of the ligand-bound PPARδ–PLIN2–CPT1A ternary complex, with enlarged views of the predicted PPARδ–CPT1A and PPARδ–PLIN2 interfaces. **B**, Representative immunoblot of global protein ubiquitination in control and hemoglobin (Hb)- exposed primary microglia treated with linoleic acid (LA; 1 μmol/L) or lysophosphatidylglycerol 16:0 (LPG[16:0]; 50 nmol/L). **C**, Coimmunoprecipitation of PPARδ followed by immunoblotting for p62 and LC3, with densitometric quantification normalized to immunoprecipitated PPARδ. **D**, Representative immunoblot of total cellular p62. **E**, Coimmunoprecipitation demonstrating reciprocal associations among PLIN2, CPT1A, and PPARδ in primary microglia; species- and isotype-matched IgG served as the negative control. **F**, Reciprocal coimmunoprecipitation of PLIN2, CPT1A, and PPARδ in control and Hb-exposed primary microglia treated with LA or LPG(16:0). Experiments were independently repeated 3 times; representative immunoblots are shown. Quantitative data are mean±SD and were analyzed using 1-way ANOVA followed by Tukey multiple-comparisons tests; *P<0.05. Band intensities were quantified using Fiji/ImageJ version 1.54p. CPT1A indicates carnitine palmitoyltransferase 1A; Hb, hemoglobin; IgG, immunoglobulin G; LA, linoleic acid; LC3, microtubule-associated protein 1 light chain 3; LPG(16:0), lysophosphatidylglycerol 16:0; PLIN2, perilipin 2; and PPARδ, peroxisome proliferator- activated receptor δ.

### The LA–LPG–PPARδ axis restores mitochondrial function by promoting lipid droplet– mitochondria coupling

To determine how mitochondria-associated PPARδ contributes to the metabolic protective effects of the LA–LPG axis, we next focused on downstream effectors involved in lipid droplet–mitochondria communication. Transcriptomic analysis showed that PLIN2 and CPT1A, two key molecules related to lipid storage and fatty acid oxidation, were enriched among genes commonly upregulated by LA and LPG (Fig. 4a). Consistently, western blot analysis confirmed that both LA and LPG significantly increased PLIN2 and CPT1A protein expression after Hb stimulation (Fig. 4b; sFig. 4a).

PLIN2 is a major lipid droplet-associated perilipin that contributes to lipid droplet stabilization and lipid storage. CPT1A is a rate-limiting outer mitochondrial membrane enzyme that converts long-chain fatty acyl-CoA into acylcarnitine, thereby enabling mitochondrial fatty acid β-oxidation. The complementary localization and functions of PLIN2 and CPT1A therefore raised the possibility that they participate in lipid droplet– mitochondria coupling during intracellular fatty acid handling.

Immunofluorescence analysis showed that Hb stimulation induced marked intracellular lipid droplet accumulation, consistent with excessive lipid storage and impaired lipid clearance. Hb also disrupted the spatial association between lipid droplets and mitochondria (Fig. 7e; sFig6b,c), suggesting inefficient transfer of accumulated lipids into mitochondria for oxidative metabolism. In contrast, LA and LPG restored lipid droplet–mitochondria coupling, accompanied by reduced lipid accumulation and improved mitochondrial metabolic activity (Fig. 7b,e).

Co-immunoprecipitation and immunofluorescence further showed that the LA–LPG axis promoted the association between PLIN2-positive lipid droplets and CPT1A-positive mitochondria in a manner linked to mitochondria-associated PPARδ (Fig. 6e,f; Fig. 7d). PPARδ was positioned to mediate the interaction between PLIN2 and CPT1A, suggesting a scaffold-like role for mitochondria-associated PPARδ in lipid droplet–mitochondria coupling. Additional co-immunoprecipitation experiments detected reciprocal interactions among PLIN2, CPT1A and PPARδ, supporting the possibility that these three proteins exist within the same molecular complex (Fig. 6e,f). Molecular docking analysis further supported a model in which PPARδ facilitates the PLIN2–CPT1A interaction (Fig. 6a).

Functionally, Seahorse analysis showed that LA and LPG restored Hb-impaired mitochondrial oxidative phosphorylation. This protective effect was abolished by etomoxir-mediated inhibition of fatty acid β-oxidation, indicating that LA–LPG-dependent mitochondrial recovery requires fatty acid oxidation. In parallel, flow cytometry showed that LA and LPG reduced Hb-induced lipid peroxidation, consistent with improved lipid handling and decreased oxidative lipid damage (Fig. 7b,f; sFig. 6a). Moreover, LA improved the expression of core electron transport chain subunits and the enzymatic activities of mitochondrial respiratory complexes (Fig. 7a,c), providing additional evidence that LA restores mitochondrial function at multiple levels.

Together, these findings indicate that LA-regulated LPG(16:0) stabilizes mitochondria-associated PPARδ, which helps maintain the PLIN2–CPT1A interaction and promotes lipid droplet–mitochondria coupling. This mechanism facilitates the clearance of Hb-induced lipid accumulation through fatty acid β-oxidation, reduces lipid peroxidation and supports mitochondrial functional recovery after Hb-induced injury.

### LA supplementation improves cognitive performance in a mouse model of subarachnoid **hemorrhage**

Behavioral outcomes were evaluated after prechiasmatic cistern blood-injection SAH in mice, as previously described with minor modifications^30,31^. To determine whether LA treatment mitigated affective behavioral abnormalities after SAH, spontaneous locomotion and center-directed exploratory behavior were assessed using the open-field test (OFT) (Fig. 8a). Mean velocity and total distance traveled were comparable among the groups, indicating that gross locomotor performance had recovered to a similar level at the time of testing and was therefore unlikely to confound the subsequent behavioral measurements (Fig. 8b). Compared with sham-operated mice, mice subjected to SAH entered the center zone less frequently and spent less time in this region (Fig. 8b). Both measures showed partial recovery in LA-treated mice relative to untreated SAH mice, indicating that LA treatment attenuated SAH-induced anxiety- and depression-like behaviors to some extent. Spatial working memory was further assessed using the spontaneous alternation rate in the Y-maze test. Consistent with the locomotor measurements described above, mean velocity and total distance traveled were comparable among the groups, indicating that differences in gross motor activity were unlikely to confound task performance (Fig. 8f). SAH reduced the spontaneous alternation rate relative to the sham group, whereas LA treatment partially restored this reduction (Fig. 8f). These findings indicate that LA treatment partially ameliorated SAH-induced spatial memory impairment. The Morris water maze (MWM) was subsequently used to assess spatial learning and reference memory (Fig. 8C–E). During the 4- day acquisition phase, escape latency progressively decreased in all groups. Compared with sham-operated mice, SAH mice exhibited a slower reduction in escape latency, whereas LA- treated SAH mice showed an improved learning trajectory, with escape latency approaching sham levels by the end of training (Fig. 8D). Average swimming speed was comparable among the groups during acquisition, suggesting that differences in escape latency were unlikely to result from impaired swimming ability. In the probe trial, total swimming distance and average speed were also comparable among the groups. However, SAH mice spent less time in the target quadrant and entered the target quadrant fewer times than sham-operated mice, indicating impaired retention of spatial memory. LA treatment increased both the time spent in and the number of entries into the target quadrant relative to untreated SAH mice (Fig. 8E). Together, the Y-maze and MWM results indicate that LA treatment partially ameliorated SAH-induced deficits in spatial working and reference memory.

**Figure 7.**
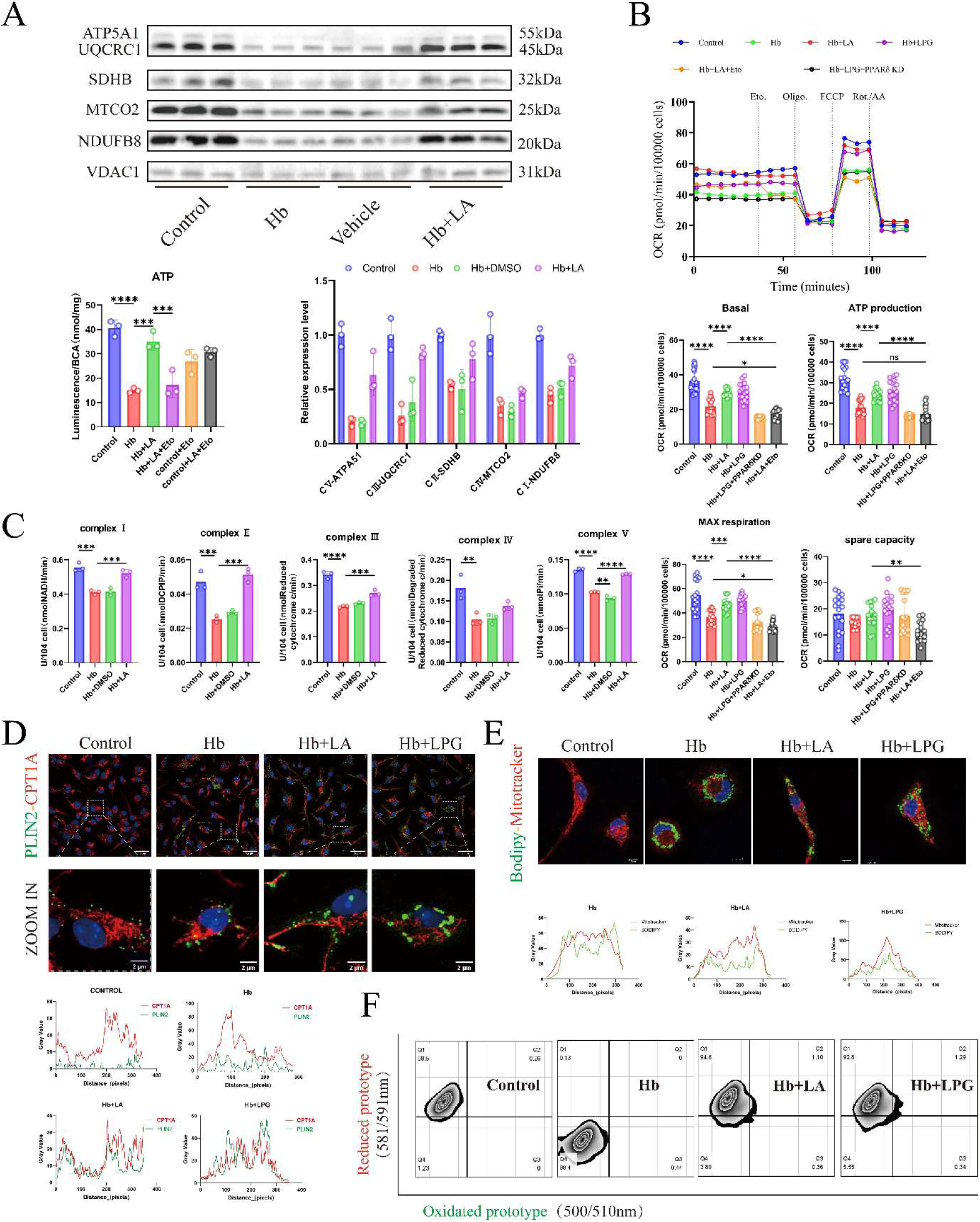
The linoleic acid–lysophosphatidylglycerol–PPARδ axis promotes lipid droplet–mitochondria coupling and mitochondrial oxidative metabolism. Primary mouse microglia were exposed to hemoglobin (Hb; 25 μmol/L) for 24 hours and treated with linoleic acid (LA; 1 μmol/L) or lysophosphatidylglycerol 16:0 (LPG[16:0]; 50 nmol/L), with etomoxir (5 μmol/L) or PPARδ knockdown as indicated. **A**, Representative immunoblots and densitometric quantification of electron-transport-chain subunits ATP5A1, UQCRC1, SDHB, MTCO2, and NDUFB8, together with intracellular ATP abundance under the indicated vehicle, LA, and etomoxir conditions. VDAC1 served as the mitochondrial loading control. **B**, Seahorse XF oxygen-consumption-rate traces and derived basal respiration, ATP-linked respiration, maximal respiration, and spare respiratory capacity. Oligomycin (1 μmol/L), FCCP (1 μmol/L), and rotenone/antimycin A (0.5 μmol/L) were injected at the indicated times. Seahorse measurements were performed in 3 independent experiments, with 5 technical replicate wells per condition in each experiment. Technical replicate wells were averaged within each experiment to generate 1 value per condition for inferential analysis (n=3 independent experiments); individual plotted points represent technical replicate wells. **C**, Enzymatic activities of mitochondrial respiratory-chain complexes I through V. **D**, Representative immunofluorescence images and line-scan profiles of PLIN2 and CPT1A. **E**, Representative BODIPY 493/503 and MitoTracker Red images and line-scan profiles showing lipid droplet–mitochondria apposition. **F**, Representative flow- cytometry plots of BODIPY 581/591 C11 oxidation as an index of lipid peroxidation. Scale bars are shown in the images. Except for the technical replicate wells displayed in B, data are mean±SD from 3 independent cultures; for B, summary statistics and inferential analyses were based on the 3 independent-experiment means. Comparisons were performed using 1- way ANOVA followed by Tukey multiple-comparisons tests; ns indicates P≥0.05; *P<0.05, **P<0.01, ***P<0.001, and ****P<0.0001. Seahorse traces were analyzed using Wave Pro version 10.2.1.4; microscopy and immunoblot signals were processed using LAS X version 3.7.22383.2 and Fiji/ImageJ version 1.54p; flow-cytometry data were analyzed using FlowJo version 10.10.0. ATP indicates adenosine triphosphate; ATP5A1, ATP synthase F1 subunit α; CPT1A, carnitine palmitoyltransferase 1A; FCCP, carbonyl cyanide-p- trifluoromethoxyphenylhydrazone; Hb, hemoglobin; LA, linoleic acid; LPG(16:0), lysophosphatidylglycerol 16:0; MTCO2, mitochondrially encoded cytochrome c oxidase II; NDUFB8, NADH:ubiquinone oxidoreductase subunit B8; OCR, oxygen-consumption rate; PLIN2, perilipin 2; PPARδ, peroxisome proliferator-activated receptor δ; SDHB, succinate dehydrogenase complex iron-sulfur subunit B; UQCRC1, ubiquinol-cytochrome c reductase core protein 1; and VDAC1, voltage-dependent anion channel 1.

**Figure 8.**
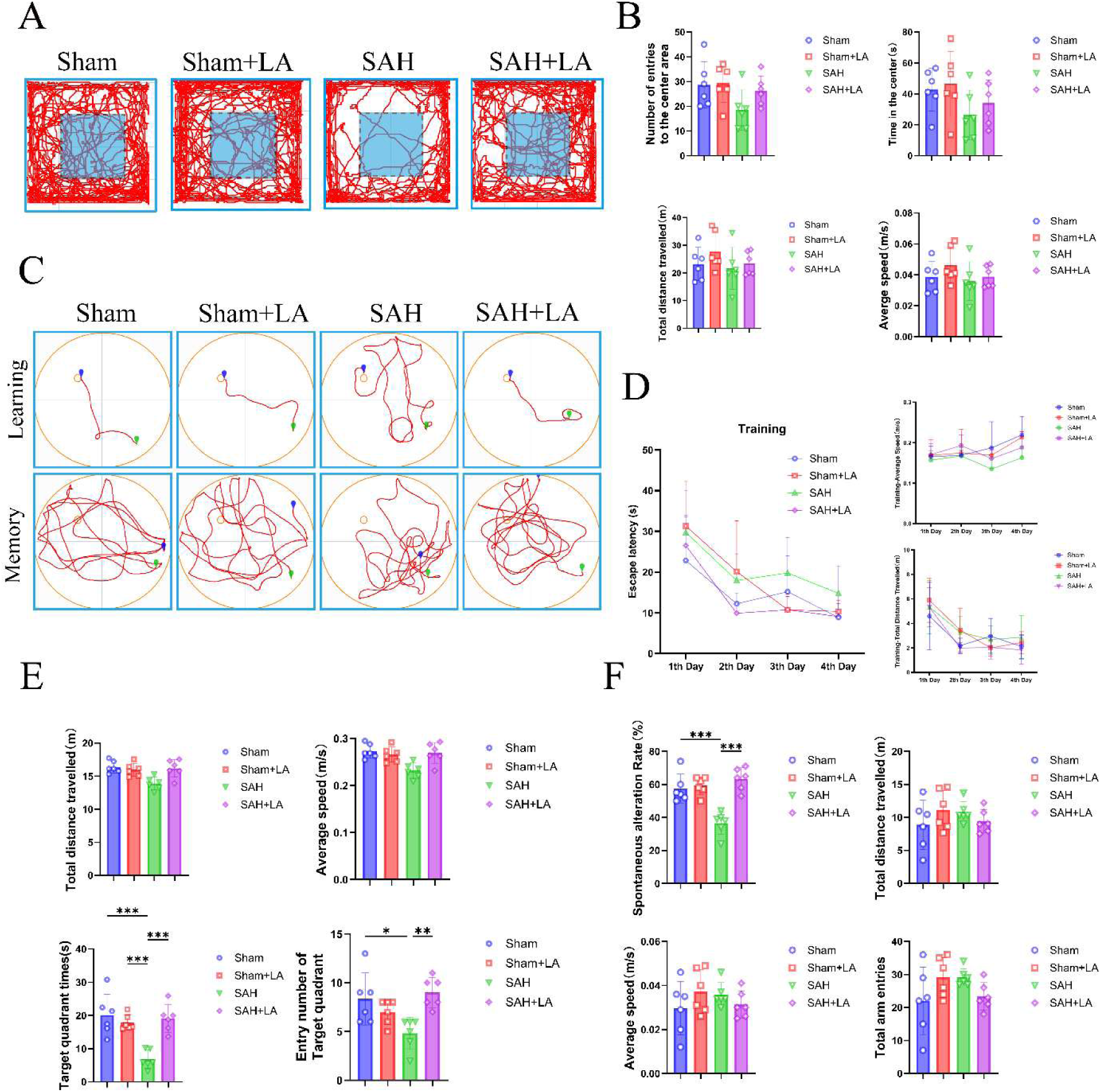
Linoleic acid improves spatial learning and memory after subarachnoid hemorrhage. Behavioral assessments were performed in sham, sham+linoleic acid (LA), subarachnoid hemorrhage (SAH), and SAH+LA mice (n=6 per group). A, Representative movement trajectories during the 5-minute open-field test; the shaded square denotes the center area. B, Number of entries into the center area, time spent in the center, total distance traveled, and average speed during the open-field test. C, Representative swimming trajectories during the learning and memory phases of the Morris water maze. D, Escape latency, training-phase average speed, and training-phase total distance traveled over 4 acquisition days. E, Total distance traveled and average speed during the probe trial, together with time spent in the target quadrant and number of entries into the target quadrant. F, Spontaneous alternation rate, total distance traveled, average speed, and total arm entries during the 5-minute Y-maze test. Behavioral testing was performed 14 days after SAH for the open-field and Y-maze tests; Morris water-maze testing began on day 16 after SAH. Data are mean±SD. The longitudinal training data in D were analyzed using 2-way repeated-measures ANOVA with treatment group and training day as factors, followed by Tukey multiple-comparisons tests. End points in B, E, and F were analyzed using 1-way ANOVA followed by Tukey multiple-comparisons tests; *P<0.05, **P<0.01, and ***P<0.001. Behavioral tracking and analysis were performed using ANY-maze version 7.67. LA indicates linoleic acid; SAH, subarachnoid hemorrhage.

## Discussion

In this translational study, we identified an LA–LPG(16:0)–PPARδ pathway linking altered lipid metabolism to microglial mitochondrial dysfunction after SAH. LA-associated metabolism was prominently altered in patient cerebrospinal fluid (CSF), whereas hemoglobin exposure reduced PLA2G15 and LPG(16:0) in primary microglia. LA restored this metabolic pathway, and LPG(16:0) preserved a mitochondria-associated pool of PPARδ. PPARδ associated with PLIN2 and CPT1A, supporting lipid droplet–mitochondria coupling, fatty acid oxidation, and respiratory recovery. These metabolic effects coincided with reduced inflammatory activation, improved phagocytosis, and partial behavioral recovery. Our findings therefore extend the metabolic interpretation of SAH from altered lipid abundance to impaired intracellular lipid utilization.

The clinical and cellular findings reveal a compartmental mismatch in LA metabolism after SAH. Although CSF LA was elevated, intracellular LPG(16:0) was markedly depleted in injured microglia. Extracellular lipid accumulation may therefore coexist with impaired routing of LA into a functional glycerophospholipid pathway. Previous metabolomic studies have similarly identified extensive changes in fatty acid mobilization and CSF lipid composition after SAH. Our findings extend these observations by connecting a clinical metabolic signal to a defined pathway in microglia. In this framework, elevated CSF LA reflects disturbed lipid homeostasis, whereas LPG(16:0) depletion marks defective intracellular lipid remodeling. The strong discrimination observed in our cohort supports LA metabolism as a candidate feature of SAH, although its biomarker performance requires validation in larger independent cohorts.

This pathway provides a metabolic explanation for several features of microglial dysfunction after hemorrhage. Hemoglobin and other blood-derived products induce inflammation, oxidative stress, mitochondrial injury, and impaired clearance of damaged cells. Previous studies have linked mitochondrial fragmentation and respiratory activity to defective phagocytosis and sustained inflammatory activation in microglia. LA and LPG(16:0) restored membrane potential, oxidative respiration, respiratory reserve, and phagocytosis while reducing oxidative and inflammatory signals. This coordinated recovery suggests that mitochondrial metabolism actively shapes microglial function rather than merely reflecting cellular injury. It also helps reconcile the apparently contrasting roles of lipid droplets in microglia. Pathological lipid droplet accumulation has been associated with oxidative stress, inflammation, and defective phagocytosis in aging and neurodegeneration. However, lipid droplets can also supply fatty acids for mitochondrial oxidation. Their functional effect may therefore depend less on abundance than on whether stored lipids remain metabolically accessible. The concurrent increases in organelle apposition, fatty acid oxidation, and respiration suggest that LA and LPG(16:0) promote productive lipid utilization rather than lipid sequestration.

LPG(16:0) emerged as the central intermediate connecting LA-responsive lipid remodeling to microglial recovery. Hemoglobin reduced PLA2G15 abundance, whereas LA restored PLA2G15 with comparatively little effect on LPGAT1. This pattern implicates impaired lysophospholipid generation, rather than generalized phospholipid acylation, in LPG(16:0) depletion. Exogenous LPG(16:0) reproduced the anti-inflammatory, pro-phagocytic, and mitochondrial effects of LA, placing it functionally downstream of the LA response. Because LA contains an 18:2 acyl chain and LPG(16:0) contains a 16:0 chain, their relationship is more consistent with LA-regulated lipid remodeling than direct conversion. PPARδ provides the next mechanistic link between this lipid signal and microglial function. Previous studies established PPARδ as a regulator of macrophage lipid metabolism, oxidative programs, and immune responses. Our data extend this framework by linking LPG(16:0) to PPARδ protein stability. Thermal stabilization, molecular simulation, and degradation experiments collectively suggest that LPG(16:0) sustains PPARδ signaling partly by limiting injury- induced protein loss.

The mitochondrial association of PPARδ adds a noncanonical dimension to this mechanism. Super-resolution imaging, subcellular fractionation, and protease protection identified a PPARδ pool accessible from the cytosolic face of the outer mitochondrial membrane.

Coimmunoprecipitation and imaging further placed PPARδ in a complex containing PLIN2 and CPT1A. Organelle contact-site studies have shown that lipid droplet–mitochondria proximity can facilitate fatty acid transfer and oxidation. However, physical proximity alone does not establish lipid flux, because increased contact can also accompany defective lipid transport. In our study, increased apposition occurred together with respiratory recovery, whereas etomoxir attenuated the metabolic rescue. These findings support functional coupling between lipid droplets and mitochondria rather than a purely morphological association. Mitochondria-associated PPARδ may therefore coordinate PLIN2-dependent lipid management with CPT1A-mediated fatty acid entry and mitochondrial oxidation. This model also explains how metabolic recovery could simultaneously reduce lipid-peroxidation stress and support energy-demanding functions such as phagocytosis. Accordingly, the observed marker changes are better interpreted as restoration of microglial metabolic flexibility than conversion to a fixed M1 or M2 phenotype.

The in vivo findings connect this cellular mechanism to neurological recovery. LA reduced molecular indices of neuroinflammation and neurovascular injury and improved spontaneous alternation, center-directed exploration, and water-maze performance. Gross locomotion and swimming speed remained comparable between groups, reducing the likelihood that motor differences explained the cognitive outcomes. Nevertheless, LA was administered before hemorrhage, so these experiments establish prophylactic biological efficacy rather than post- SAH therapeutic effectiveness. Other limitations also define the scope of the findings. The clinical cohort was small, single-center, and hospital-based, unmatched case-control study, and controls were selected according to clinical indications for CSF sampling. The direct interaction between LPG(16:0) and PPARδ, the responsible binding residues, and the necessity of the PLA2G15–LPG(16:0) pathway require biochemical and genetic validation. Isotope-resolved flux analysis is also needed to measure lipid transfer between droplets and mitochondria directly. Finally, studies incorporating female animals, microglia-specific PPARδ deletion, and post-hemorrhage treatment will be important for assessing generalizability and translational feasibility.

In conclusion, LA restores PLA2G15-associated LPG(16:0) signaling and preserves mitochondria-associated PPARδ after SAH. Through its association with PLIN2 and CPT1A, PPARδ links lipid remodeling to productive lipid droplet–mitochondria coupling, fatty acid oxidation, and microglial recovery. The LA–LPG(16:0)–PPARδ axis therefore provides a mechanistic framework for post-hemorrhagic immune dysfunction and a candidate target for further translational investigation.

## Acknowledgments

The authors thank the patients and their families for participating in this study. We also thank the clinical staff for assistance with cerebrospinal fluid collection and clinical data management and the institutional metabolomics and imaging core facilities for technical support. An OpenAI large language model was used to assist with English translation and language editing. All generated text was critically reviewed and revised by the authors, who take full responsibility for the final content.

## Sources of Funding

This work was supported by the National Natural Science Foundation of China (NO.U24A20687, NO.82130037 for C.H.; NO.82401551 for Tao.Tao.) and the Natural Science Foundation of Jiangsu Province (No. BK20240016 for W. L.)

## Disclosures

None.

## Data Access and Responsibility

Le-Xuan Zou had full access to all data in the study and takes responsibility for the integrity of the data and the accuracy of the data analysis.

## Supplemental Material

Supplemental Methods

Supplemental Figures 1–6 Major Resources Table

Supplemental Table 1. Patient Baseline Characteristics

Supplemental Table 2. Animal Allocation, Exclusions, and Mortality

## Non-standard Abbreviations and Acronyms

AUC,: area under the curve;
CPT1A,: carnitine palmitoyltransferase 1A;
CSF,: cerebrospinal fluid; Hb, hemoglobin;
LA,: linoleic acid;
LPG(16:0),: lysophosphatidylglycerol 16:0;
OCR,: oxygen-consumption rate;
PLIN2,: perilipin 2;
PPARδ,: peroxisome proliferator-activated receptor-δ;
SAH,: subarachnoid hemorrhage.

## Notes

### Competing Interest Statement

The authors have declared no competing interest.

